# Climate change alters plant-herbivore interactions

**DOI:** 10.1101/2020.08.31.275164

**Authors:** Elena Hamann, Cameron Blevins, Steven J. Franks, M. Inam Jameel, Jill T. Anderson

## Abstract

Plant-herbivore interactions have evolved in response to co-evolutionary dynamics, along with selection driven by abiotic conditions. We examine how abiotic factors influence trait expression in both plants and herbivores to evaluate how climate change will alter this long-standing interaction. The paleontological record documents increased herbivory during periods of global warming in the deep past. In phylogenetically-corrected meta-analyses, we find that elevated temperatures, CO_2_ concentration, drought stress and nutrient conditions directly and indirectly induce greater herbivore consumption, primarily in agricultural systems. Additionally, elevated CO_2_ delays herbivore development, but increased temperatures accelerate development. For annual plants, higher temperatures, CO_2_, and drought stress increase foliar herbivory, and our meta-analysis suggests that greater temperatures and drought may heighten florivory in perennials. Human actions are causing concurrent shifts in CO_2_, temperature, precipitation regimes and nitrogen deposition, yet few studies evaluate interactions among these changing conditions. We call for additional multifactorial studies that simultaneously manipulate multiple climatic factors, which will enable us to generate more robust predictions of how climate change could disrupt plant-herbivore interactions. Finally, we consider how shifts in insect and plant phenology and distribution patterns could lead to ecological mismatches, and how these changes may drive future adaptation and coevolution between interacting species.

## INTRODUCTION

Plant-herbivore interactions structure the ecology of natural communities and evolutionary trajectories of interacting species (Becerra, 2007). Herbivores first began to consume plant tissues in the late Silurian to early Devonian (e.g., Edwards *et al*., 1995; Labandeira & Currano, 2013), shortly after land plants evolved approximately 475 million years ago (Wellman *et al*., 2003). Since then, plants have evolved complex mechanical and chemical defenses against herbivory (Gong & Zhang, 2014; Burkepile & Parker, 2017), and herbivores have evolved to circumvent these defenses (Karageorgi *et al*., 2019). Coevolutionary dynamics can generate specialized interactions, resulting in a diversity of species, forms and functional traits (Jander, 2014; Karageorgi *et al*., 2019). However, even intense and prolonged eco-evolutionary dynamics occur in the context of an abiotic environment that can change. Given that the abiotic environment strongly influences trait expression of both plants and herbivores, changes in abiotic conditions can alter this ecologically-important interaction. For example, during the intense planetary warming and increased CO_2_ levels of 55.8 million years ago, fossilized plants show clear evidence of exposure to a greater degree and diversity of herbivory than in cooler times (Currano *et al*., 2008; Pinheiro *et al*., 2016).

In contemporary landscapes, natural populations are encountering novel suites of abiotic and biotic conditions as a result of rising atmospheric CO_2_ concentrations, global temperatures, changing precipitation regimes, and increased nitrogen deposition (Bellard *et al*., 2012; IPCC, 2014). Climate-change-mediated shifts in plant-herbivore interactions could reshape the evolutionary trajectories and ecological dynamics of entire communities (Rasmann *et al*., 2014b; Becklin *et al*., 2016). Insect herbivores have short generation times, high reproductive rates, and extensive mobility patterns (Menéndez, 2007). All of these traits could enable them to track favorable climates more readily than plants, through rapid migration, *in situ* population growth, and adaptation to novel suites of climatic conditions (Rasmann *et al*., 2014a; Becklin *et al*., 2016), which could lead to non-analog biotic assemblages (Parmesan, 2006). For these reasons, plant populations could experience increased levels of herbivory under continued climate change.

Here, we synthesize existing literature and conduct meta-analyses to evaluate the extent to which rapid anthropogenic climate change disrupts plant-herbivore interactions. Specifically, we hypothesize that climate change will expose native plants to: (1) novel herbivore communities both in their home sites and expanded distributions; and (2) heightened levels of herbivory because of increased food consumption by resident and newly-established herbivores. We first review the proximate mechanisms by which warming temperatures in concert with elevated CO_2_, novel precipitation patterns, and increased nitrogen deposition influence herbivore biology and plant damage from herbivory. We consider how multiple interacting climatic factors could result in additive, synergistic or antagonistic effects for plants-herbivore interactions. By reflecting on spatial and temporal shifts in plant-herbivore interactions, we explore whether natural plant populations will confront novel patterns of herbivory under future climates. Our review primarily focuses on arthropods (Hexapoda), which represent about ∼62% of all ∼1.6 million described living species, as approximately half of all insects are herbivorous (Wiens *et al*., 2015; Roskov *et al*., 2019). Insects have received more attention in the context of plant-herbivore interactions under climate change, yet we draw on mammalian herbivore examples when possible.

To complement the qualitative literature review, we test whether climate change will alter herbivore biology and augment herbivory in natural and agricultural systems in a series of meta-analyses. To that end, we ask how abiotic factors associated with climate change affect herbivore performance and feeding rates. We consider whether climate change factors affect herbivores directly or indirectly via plant-mediated effects, and examine potential differences across study systems. We then investigate herbivore damage records under simulated climate change to evaluate how plant populations – adapted to historical levels of herbivory – fare under novel herbivory associated with climate change.

Several earlier reviews and meta-analyses concentrated on the effects of elevated CO_2_ on plant chemistry and functional traits, and herbivore performance (Bezemer & Jones, 1998; Zvereva & Kozlov, 2006; Stiling & Cornelissen, 2007; Robinson *et al*., 2012). Additionally, Robinson *et al*. (2012) examined pairwise interactions between CO_2_ and temperature, CO_2_ and drought, and CO_2_ and nitrogen for plant phenotypes, but they did not consider the effects of these interactions on insect performance or plant damage from herbivores. As changes in plant chemistry and phenotypes have been established in response to elevated CO_2_ and temperature, we did not assess such responses. However, only 10 empirical studies included in previous meta-analyses examined herbivore damage to plants under climate change (Stiling & Cornelissen, 2007; Robinson *et al*., 2012). Thus, we quantified herbivore responses and plant damage to multiple climate change factors across empirical studies. Moreover, while some previous studies considered the effects of feeding guilds or insect orders in their meta-analyses (Bezemer & Jones, 1998; Stiling & Cornelissen, 2007; Robinson *et al*., 2012), previous efforts did not account for phylogenetic relatedness, which could have generated anti-conservative results. Finally, we fill a gap in the literature by examining differences in climate change responses in native vs. agricultural systems.

After evaluating how climate change factors influence plant-herbivore dynamics in the short-term, we discuss how these proximate causes may ultimately alter plant-herbivore interactions in light of evolutionary dynamics and examine the potential for plants and their herbivores to adapt to novel abiotic and biotic pressures.

## I. Literature Review: Proximate ecological responses of plants and herbivores to climate change

### Elevated atmospheric CO_2_

Atmospheric CO_2_ concentrations have risen from 280 ppm during pre-industrial times to the current 410 ppm, and are predicted to exceed 600 ppm by the end of the 21^st^ century (NOAA, 2020). Although elevated CO_2_ has little direct effects on insect herbivores (Kerr *et al*., 2013), it can indirectly influence herbivores via plant-mediated effects (Pincebourde *et al*., 2017). Increased atmospheric CO_2_ levels alter the carbon (C) and nitrogen (N) economy within the plant (increased C:N ratio), decreasing the N levels in plant tissue (Strain, 1987; Fajer, 1989; Johnson & Lincoln, 1990). As N is a limiting nutrient for insects (Mattson 1980), higher CO_2_ levels diminish the nutritional quality of plant tissues by reducing concentrations of proteins and certain amino acids in leaves (Lincoln *et al*., 1993; Docherty *et al*., 1997). To compensate, insect herbivores can increase their food uptake (Johnson & Lincoln, 1990; Johnson & Lincoln, 1991; Stiling & Cornelissen, 2007). Decreased foliar N content and increased defenses can reduce the conversion efficiency of ingested food (Fig. 1). The most extensive meta-analysis on herbivore responses to elevated CO_2_ confirmed these patterns by examining 270 papers published between 1979 and 2009 (Robinson *et al*., 2012). In response to a 19% increase in foliar C:N and a 10% decline in foliar proteins, insect herbivores increased their relative consumption rates by 14%, yet the conversion efficiency decreased by 15% under elevated CO_2_ (Robinson *et al*., 2012). However, in line with an earlier study showing that only leaf-chewers significantly increased their food uptake in elevated CO_2_ conditions (Bezemer & Jones, 1998), Robinson *et al*. (2012) found that consumption rates increased mainly in foliage-feeders, particularly in Lepidoptera and Coleoptera. A taxonomic bias toward Lepidopteran herbivores (Bezemer & Jones, 1998; Stiling & Cornelissen, 2007) may drive the assumption that herbivore consumption will increase under climate change, although this may not be universally true across herbivore orders and feeding guilds.

**Fig. 1:**
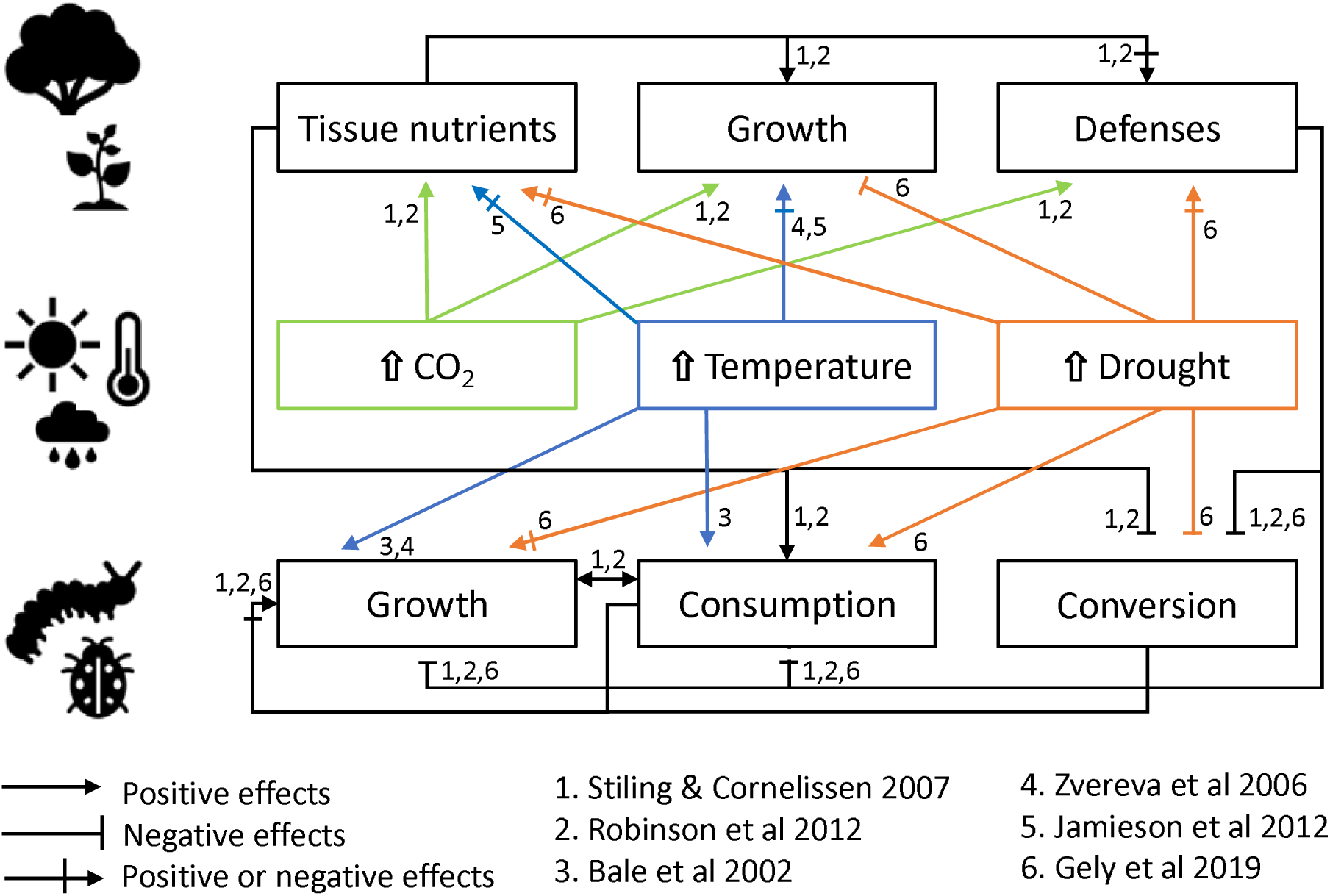
Effects of single climate change factors (in the middle) on plant growth, tissue nutrients status and defenses (top), and insect growth, consumption, and food conversion efficiency (bottom). Lines ending with arrowheads represent positive effects, lines ending in a vertical bar represent negative effects and lines with both symbols represent either positive or negative effects. We supplemented the represented effects with references from reviews or quantitative meta-analyses (represented by numbers).

Plants do not necessarily suffer more damage even when per capita consumption rates increase, since plants can accumulate more biomass under elevated CO_2_ (Hunter, 2001; Reddy *et al*., 2010). Thus, enhanced plant growth from elevated CO_2_ could compensate for increased leaf damage (Hughes & Bazzaz, 1997; Hall *et al*., 2005). However, multiple climatic factors are changing simultaneously, and increased herbivore population growth rates induced by warming temperatures could compound overall rates of herbivory. Increased rates of herbivory could impose strong selection on plant populations for constitutive and inducible defenses against herbivory. Future studies can evaluate evolutionary responses by quantifying genetic variation in defenses within- and across-populations in conjunction with estimating gene flow rates, comparing patterns of selection on defenses under historical, contemporary and future climates, and taking resurrection approaches to test directly for adaptive responses to climate change and increased herbivory (see Section 3 on evolutionary response) (Agrawal *et al*., 2006; Franks *et al*., 2018a).

### Rising temperatures

Greenhouse gas emissions have increased global temperatures by 1.0 ± 0.2 °C since 1880, and temperatures are projected to rise 2-4°C relative to preindustrial climates by 2100 under current rates of climate change (IPCC, 2018). Temperature regulates the metabolism and physiology of ectotherms, and directly influences all components of the life history of arthropod herbivores (Bale *et al*., 2002). Accelerated metabolic rates of herbivores under elevated temperature could lead to higher consumption, growth and faster development (Fig. 1), which would increase population growth rates and reduce generation times (Cornelissen, 2011; Jamieson *et al*., 2012). Furthermore, warmer winters and earlier springs associated with climate change could increase herbivore overwinter survival (Bale *et al*., 2002). Warming-induced insect outbreaks could become more frequent (Coley, 1998). For example, recent warming has caused the mountain pine beetle (Dendroctonus *ponderoae*) to shift from a semivoltine to a univoltine lifecycle, and larger herbivore outbreaks significantly increased damage to whitebark pine trees (*Pinus albicaulis*) in Yellowstone National Park (Logan *et al*., 2010). These rapid changes in herbivore performance and life-history traits could render plant populations more vulnerable to herbivores.

While increased temperatures tend to have positive effects on insects (Bale *et al*., 2002; Cornelissen, 2011), the biological impacts of rising temperatures depend on the magnitude of the change and on the herbivore’s thermal sensitivity. Insect performance typically increases with temperature until reaching a maximum at an intermediate temperature and then rapidly decreasing (Kingsolver, 2009). The asymmetry in thermal performance curves could result in very different short-term responses to increased temperatures, depending on the current location along the curve (Fig. 2). Additionally, projected climate warming could augment insect performance at temperate and poleward latitudes, where species have broader thermal tolerance, but have deleterious consequences for tropical species with more narrow thermal tolerances where temperatures may already be close to optimal (Deutsch *et al*., 2008; Angilletta *et al*., 2010). In the longer-term, rapid generation times and intraspecific genetic variation in heat resistance may allow for rapid adaptation of thermal performance curves as temperatures continue to increase (Muñoz-Valencia *et al*., 2016; Ranga *et al*., 2017). For example, Carbonell and Stoks (2020) recently documented evolutionary changes in the thermal performance curves of the European damselfly (*Ischnura elegans*) during its range expansion toward warmer and cooler regions. Similar studies examining thermal performance in herbivores are needed, as it remains unclear whether standing levels of genetic variation for heat resistance are adequate for sustained responses to selection, which may rapidly plateau (Kellermann *et al*., 2012; Kellermann & van Heerwaarden, 2019).

**Fig. 2:**
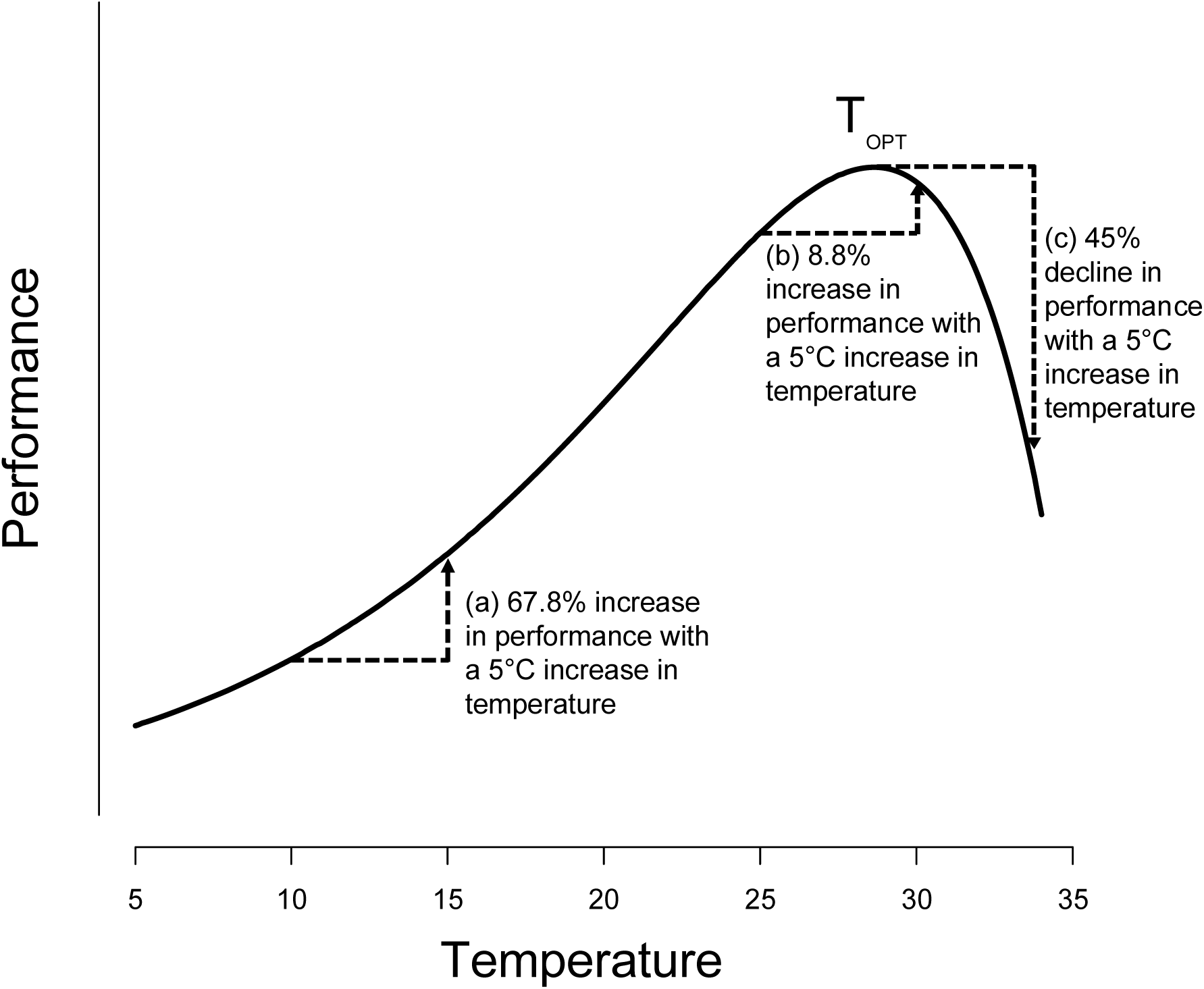
A hypothetical thermal performance curve for an insect herbivore with optimal performance (T_opt_) at 28.6 °C. Due to the asymmetry of the curve, an increase in temperature by 5°C leads to a much larger rise in performance (a) at low temperatures than (b) when the temperature approaches the optimum. As temperatures warm above the optimum, insect performance declines precipitously (c), as shown by a 45% reduction in performance under temperatures 5°C greater than the optimum. We simulated this hypothetical curve using the R package thermPerf (ver. 0.0.1) (Bruneaux, 2016).

Climate change can also affect the number of life cycles that can be completed in a single season (voltinism). Insect species typically respond to warmer temperatures with faster developmental rates and greater reproductive potential, which can increase both the number of generations within a season and the rate of population growth (Jönsson *et al*., 2009; Altermatt, 2010; Fand *et al*., 2014). These changes could intensify herbivory pressures and increase levels of damage to plants, especially to long-lived species (DeLucia *et al*., 2012; Forrest, 2016). Yet, climate change will not induce advanced emergence or voltinism in all species (Grevstad & Coop, 2015), especially those that have obligate diapause with required chilling periods that may be prolonged under warmer winters (Harrington *et al*., 1999; Forrest, 2016). Additionally, rapid development at higher temperatures is often accompanied by smaller size at maturity and reduced fecundity (Kingsolver & Huey, 2008). Yet, accelerated population growth rates increases the general abundance of insect herbivores, regardless of changes in voltinism and emergence times, and may increase herbivory on host plants.

Temperature directly influences plant growth and development (Grace, 1987). A moderate rise in temperature can increase plant productivity and production of secondary metabolites (Rustad *et al*., 2001; Zvereva & Kozlov, 2006; Pincebourde *et al*., 2017). However, severe drought could accompany heat waves associated with climate change and have adverse effects on plant productivity (Fig. 1). Plant populations located in regions that historically experienced low levels of herbivory could be susceptible to decline if herbivore populations expand *in situ* and migrate into those areas. Biotic interactions, including herbivory, have historically been considered less intense at poleward latitudes, where plants have in turn evolved fewer defense traits (Coley & Aide, 1991; but see Moles *et al*., 2011); thus, plants could be particularly susceptible to increased herbivore pressures in those regions. It thus remains an open question whether plants will suffer greater damage if herbivore performance increases or generation times decrease with rising temperatures.

### Drought

Under climate change, precipitation may increase in some parts of the world, but many other regions are projected to suffer more frequent and severe drought events (Knapp *et al*., 2008; IPCC, 2014; Swain *et al*., 2014). In combination with elevated temperatures and heat waves, drought is likely to affect many ecosystems (Jentsch *et al*., 2007; Knapp *et al*., 2008; Bloor *et al*., 2010). Additionally, in snow-dominated ecosystems at high elevations and high latitudes, climate change is causing a reduction of winter snowpack (Fyfe *et al*., 2017), which is critical as water availability to plants in the growing season can be dominated by snowmelt (Jamieson *et al*., 2012). In general, drought negatively affects plant productivity (Fig. 1), and alters plant chemical defenses and nutritional quality, digestibility and palatability (Jamieson *et al*., 2017). Traditionally, herbivorous insects were thought to perform better on water-stressed plants (Huberty & Denno, 2004). During droughts, herbivore outbreaks could result from accumulations of nitrogen compounds in plant tissue, which enhance herbivore growth and reproduction (White, 1984). However, Huberty and Denno (2004) showed that prolonged water-stress in plants negatively affects phloem and mesophyll feeders and other sap-feeders, while the responses of leaf chewers differed between sub-guilds. Although foliar nitrogen concentrations generally increase during times of water deficit, decreased turgor and water availability can interfere with the herbivore’s ability to access nitrogen, especially in continuously-stressed plants (Huberty & Denno, 2004). Intermittent and moderate droughts may increase chemical defenses in plants, while prolonged and severe droughts could decrease these defenses; in both cases, limited access to nutritional compounds could reduce herbivore performance (Gely *et al*., 2019). The effect of drought stress on chemical defense production and nutritional quality has not yet been tested as rigorously as the effects of elevated temperature or CO_2_. Additionally, species-specific plant and herbivore physiology needs to be considered to understand these interactions under drought.

### Fertilization and N deposition

Fertilization is commonly used in agricultural systems, leading to agricultural runoff and increased N deposition in many parts of the world (Hattenschwiler & Schafellner, 1999; Driscoll *et al*., 2003). Intensified N deposition can stimulate plant growth in the short-term (Tamm, 1991; Hattenschwiler & Schafellner, 1999), and increase the nutritional value of plant tissues (Henn & Schopf, 2001). However, Hattenschwiler and Schafellner (1999) found that the stimulating effects of increased N deposition were lower in magnitude than the adverse effects of elevated CO_2_, such that climate change may still impair herbivore growth and development. N deposition could allow plants to allocate more resources to herbivore defenses (plant vigor hypothesis), or could exacerbate herbivory because the stress caused by the pollution reduces investment in defense (plant stress hypothesis) (Mur *et al*., 2017; Miles *et al*., 2019). Moreover, drought severely limits the ability of plants to acquire soil nutrients (Bista *et al*., 2018). Thus, it is important to consider fertilization at a regional level and in combination with other climate change factors.

### Interactions between abiotic factors

Climate change is simultaneously altering key agents of selection, including CO_2_, temperature, and precipitation patterns, while increasing the frequency and severity of extreme weather (IPCC, 2014). In addition, habitat fragmentation restricts migration and could hinder adaptive responses to novel environments, as small fragmented populations often lack genetic variation (Young *et al*., 1996; Leimu *et al*., 2006). Global change factors interact in complex ways that could have additive, synergistic or antagonistic effects on plant-herbivore interactions (Jamieson *et al*., 2017; Gely *et al*., 2019). While elevated CO_2_ is increasing at similar rates globally, changes in temperature and precipitation patterns vary regionally and inter-annually (Jentsch *et al*., 2007; IPCC, 2014; Swain *et al*., 2018). Yet, few empirical studies explore herbivore and plant performance under realistic, multifactorial scenarios (Zvereva & Kozlov, 2006; Cornelissen, 2011). In contrast, plant physiological responses to climate change have been relatively well-studied in a multifactorial framework (e.g., Veteli *et al*., 2002; Williams *et al*., 2003; Murray *et al*., 2013). For example, elevated CO_2_ and temperature significantly increased foliar C:N ratios in two Eucalyptus species (*E. robusta and E. tereticornis*) especially under ambient temperatures (Murray *et al*., 2013; Gherlenda *et al*., 2015). In other studies, nitrogen content decreased strongly under combined elevated CO_2_ and temperature, while elevated temperatures alone had little effect (Williams *et al*., 2000; Murray *et al*., 2013; Gherlenda *et al*., 2015). In a synthesis of 42 studies, Zvereva and Kozlov (2006) confirmed that the combined effects of elevated CO_2_ and temperature on plant physiology and chemistry were often different from the effects of factors taken separately. Thus, multifactorial experiments can reveal unique biological responses to climate change that are not apparent from single factor studies (e.g., Zvereva & Kozlov 2006; Robinson et al., 2012).

Even less is known about interactive effects of climate change factors on herbivore performance than on plant physiology. In single factor studies, insect performance declines under elevated CO_2_ and increases under warmer temperatures, but when manipulated simultaneously, the effects often cancel each other out (Williams *et al*., 2000; Johns & Hughes, 2002; Veteli *et al*., 2002; Johns *et al*., 2003; Williams *et al*., 2003; Chong *et al*., 2004). More recently, Zhang et al. (2018) found that elevated CO_2_ and temperature significantly decreased growth rates and conversion efficiency of consumed food in *Spodoptera litura*, but other studies detected no significant interactions between these climate change factors (Himanen *et al*., 2009; Murray *et al*., 2013; Niziolek *et al*., 2013; Gherlenda *et al*., 2015). The stimulating effects of temperature on insect performance may not ameliorate the negative plant-mediated effects of elevated CO_2_. Yet, enhanced temperature and CO_2_ could both increase foliar damage via increased insect abundance, growth and consumption (Niziolek *et al*., 2013). We cannot make reliable predictions about plant-herbivore interactions under climate change until future studies explicitly evaluate the combined effects of climate change factors on herbivore performance, changes in host plant quality, and plant damage.

### Spatial and temporal mismatches between plants and herbivores

Climate governs the geographic distribution of many species (e.g., Sexton *et al*., 2009), and climate change has already led to shifts in species distribution (Parmesan & Yohe, 2003; Root *et al*., 2003; Thuiller, 2004; Pereira *et al*., 2010). These distributional changes include expansions into new areas, especially at the historically cooler, upper elevational and poleward latitudinal limits (leading edges), and local extinctions in areas that have become climatically unsuitable, especially at the warm lower elevational and latitudinal limits of species ranges (trailing edges) (Menéndez, 2007; Sheth & Angert, 2018). As a result, ecological communities may disassemble as individual species shift their ranges idiosyncratically, and new assemblages are likely to emerge (Thuiller, 2004; Leimu *et al*., 2012; Maron *et al*., 2019), which will likely influence plant-herbivore interactions (Harrington *et al*., 1999; Menéndez, 2007).

Plant species could confront novel herbivore communities and novel levels of herbivory both in their home sites and during distributional shifts. Herbivores may have greater migratory potential than plants, and fossil records show that many insect species tracked climate change via migration during geological periods of climate change (Coope, 1970; Lawton, 1995). As climates change, thermal limits may no longer constrain native herbivores to their historical ranges. For example, the mountain pine beetle (*D. ponderoae*), native to western North America, is currently expanding its range north-eastward establishing on novel host trees, such as jack pines (*Pinus banksiana*) as it spreads through the boreal forest (Cullingham *et al*., 2011; Rosenberger *et al*., 2017). This example illustrates how the oligophagous or polyphagous diet of many insect herbivores may facilitate host shifts during range expansion, exposing many plant species to novel herbivory, and driving host range evolution in insects (Agrawal, 2000).

Climate change is accelerating the timing of life history events for many species (Parmesan & Yohe, 2003). Yet, the environmental cues that trigger phenological transitions, and their relative importance, often differ between plants and herbivores, resulting in phenological mismatches between interacting species and trophic levels (Choi *et al*., 2019). In general, arthropod herbivores are advancing their phenology faster than plants, as they are more sensitive to temperature while plants often have specific photoperiod thresholds (Visser & Christiaan, 2005; Menéndez, 2007; Körner & Basler, 2010; but see Forrest & Thomson, 2011). Asynchronous phenological shifts may generate temporal mismatches, which may amplify or dampen herbivore damage (Dewar & Watt, 1992; Diamond *et al*., 2011; DeLucia *et al*., 2012; Abarca & Lill, 2015; Ren *et al*., 2020). Warm springs in temperate regions often induce earlier insect emergence and activity, especially for insects that overwinter as adults (Diamond *et al*., 2011; Bell *et al*., 2015). Consequently, herbivory could increase early in the season, and herbivores may have extended growing seasons (Forrest, 2016). In specialized interactions, phenological asynchrony may reduce herbivore growth and abundance if climate change causes larvae to emerge earlier than bud-burst of the host species (Visser & Holleman, 2001; Schwartzberg *et al*., 2014). Yet, many generalist herbivore species may be resilient to phenological changes in host plants (Forrest & Thomson, 2011). For example, spring herbivore species have often evolved starvation tolerance, enabling them to survive when hatching occurs before bud burst (Abarca & Lill, 2015; Kharouba *et al*., 2018). Phenological shifts can also remove temporal barriers (Kharouba *et al*., 2018). For example, warming synchronized the hatching time of forest tent caterpillar (*Malacosoma disstria*) eggs and budburst of one tree host, but reduced synchrony with an alternate host (Visser & Holleman, 2001). Starvation endurance and broad dietary breadth may dampen the effects of altered plant phenology for herbivores, but shifting temporal dynamics could augment herbivore damage on plants.

## II. META-ANALYSIS

### Aims and hypotheses

We conducted phylogenetically-corrected meta-analyses (Adams, 2008) to evaluate: (1) the direct, indirect (i.e. plant-mediated) and total effects of climate change on arthropod herbivore performance and foraging biology; and (2) the effects of climate change on plant damage, in natural and agricultural systems. We sought to test whether rapid contemporary climate change augments herbivory, similar to what occurred during periods of elevated atmospheric [CO_2_] in the geological record (Currano *et al*., 2008). We aimed to assess the interactive effects of climate change factors, but our statistical power was restricted by the low numbers of multifactorial studies (see Results and Discussion section).

### Methods

In the supplementary materials, we describe study eligibility criteria, the literature search, data extraction, phylogeny reconstructions, data analysis, and publication bias diagnostics (illustrated in Fig. S1). In short, we used Web of Science to conduct literature searches from 1900 to August 6, 2020 on (1) herbivore performance and (2) herbivory in climate manipulation studies. We also extracted data from studies cited in previous meta-analyses (Stiling & Cornelissen, 2007; Robinson *et al*., 2012) that measured herbivore performance and consumption rates in response to different CO_2_ levels. Finally, we performed a forward search from Robinson (2012) for publications that reported herbivore performance or herbivory in response to climate change. For herbivores, we concentrated on individual growth rates, development time, and consumption rates, and population-level metrics, including abundance and population growth rates. For plants, we focused on measures of tissue damage caused by herbivore feeding, such as damaged leaf area, percentage tissue loss, and feeding marks. We extracted data directly from tables, archived datasets, or figures using WebPlotDigitizer (Rohatgi, 2019) for all papers fitting our eligibility criteria (N= 62 studies for herbivore performance, 26 of which had not been included in previous meta-analyses, and N= 47 for plant damage, 33 of which were unique to this meta-analysis, Fig. S1, S4, S5, S7, S8, Table S12 and S13).

Studies in the herbivore dataset used three different experimental designs. Some studies evaluated the direct effects of climate change factors on herbivores by exposing individuals to experimental manipulations while feeding them an artificial diet or leaves of plants grown under ambient conditions. Other studies evaluated the indirect effects of climate change on herbivores mediated through plants by rearing herbivores under ambient conditions and feeding them tissue from plants exposed to climate change manipulations. Finally, the last set of studies tested the total effects of climate change by exposing both the plants and the herbivores to manipulations and monitoring herbivore responses. We present results from the full dataset and then dissect the direct, indirect and total effects of climate change through separate analyses of subsets of data.

Our datasets also included agricultural, biocontrol and native plant and herbivore species. To our knowledge, previous meta-analysis have not tested whether shifts in plant-herbivore interactions under climate change are consistent across wild and domesticated systems. However, a recent phylogenetically corrected meta-analysis showed that plant resistance to herbivores was lower in domesticated crops relative to their wild relatives (Whitehead *et al*., 2017). We present results for the full herbivore and plant datasets, and then evaluate herbivore performance and plant damage under climate change factors in agricultural and biocontrol vs. native systems.

We constructed phylogenies (Fig. S2 and S3) based on publicly-accessible data in the Open Tree of Life (Michonneau *et al*., 2016) to include a phylogenetic correlation matrix in our models. We implemented multilevel mixed-effects meta-analysis in the R package metafor, using Hedges’ *g* (Viechtbauer, 2010). We computed effect sizes such that values <0 indicated that treatments consistent with climate change projections (e.g., increased CO_2_) depressed herbivore performance or leaf damage from herbivores, and effect sizes > 0 indicated that climatic manipulations augmented herbivore performance or leaf damage from herbivores. The final models included fixed effects (moderating factors) for climatic manipulations and herbivore or plant traits and other attributes of the studies (publication year, latitude, longitude, elevation, study setting, etc.), and random effects for publication (to account for multiple species or traits in a study) and the phylogenetic correlation matrix.

### Results and Discussion

#### Herbivore dataset

Our analyses revealed significant effects of herbivore trait, climatic treatment and their interaction (Fig. 3, Table S1-S7). Across analyses, elevated CO_2_, temperature, drought stress and fertilization all increased herbivore consumption rates (Fig. 3), suggesting that herbivore pressures are intensifying under most climate change scenarios.

**Fig. 3:**
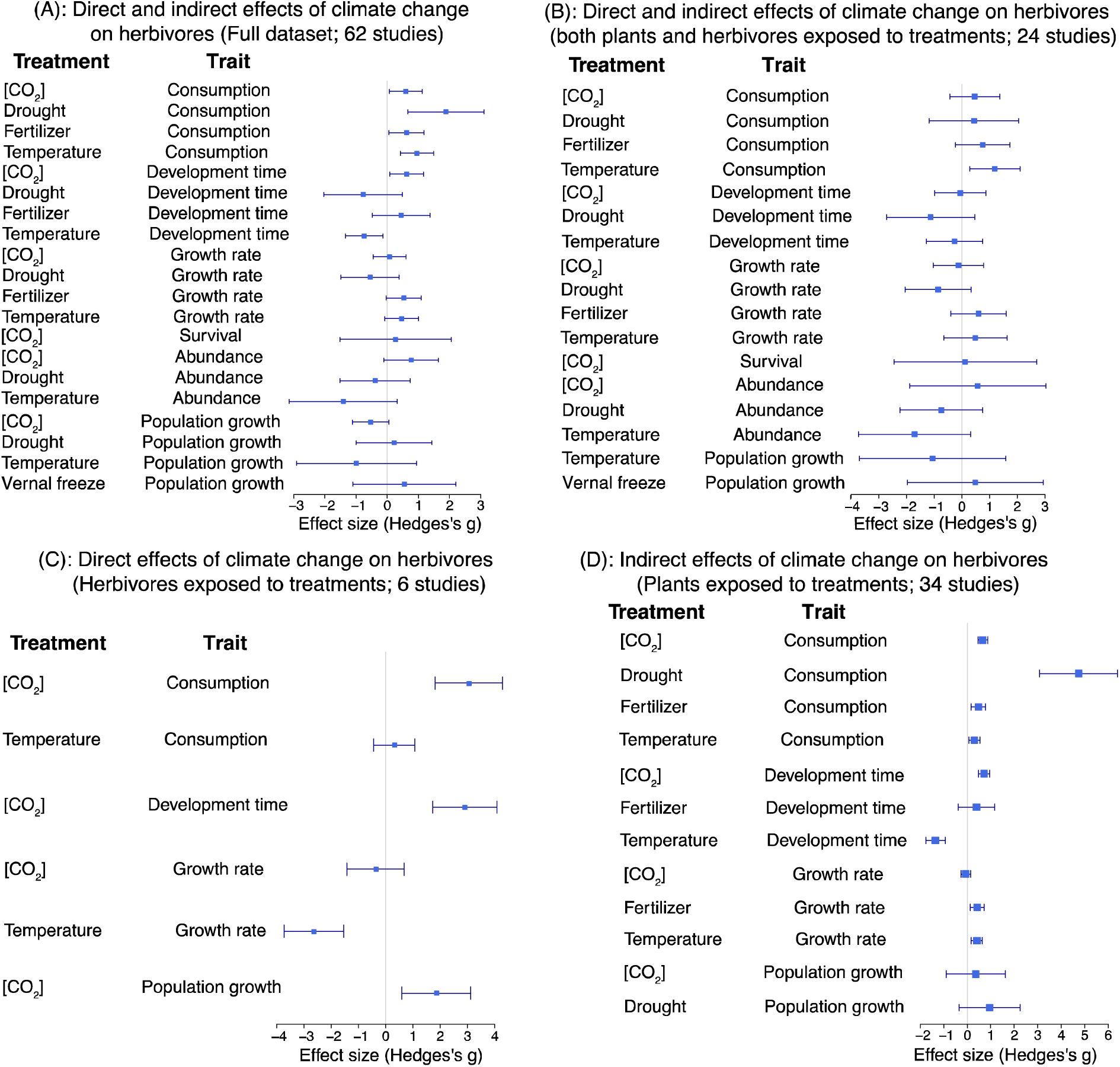
Results of phylogenetically-corrected meta-analysis of herbivore responses at the individual- and population-levels to climate change manipulations in the field and controlled conditions. Plotted are Hedges’s g effect sizes and 95% confidence intervals. (A) The full dataset includes 61 studies that varied in experimental design. This panel represents the results of all studies combined. We also conducted meta-analyses of three subsets of data, representing: (B) studies that exposed both herbivores and plants to experimental treatments, documenting both direct and indirect effects of climate change on herbivores; (C) a small number of studies that exposed herbivores (but not plants) to elevated temperatures and atmospheric [CO_2_], thus demonstrating the direct effects of climate change on herbivores; and (D) studies that exposed plants (but not herbivores) to experimental manipulations, revealing indirect effects of climate change on herbivores. Positive values with 95% confidence intervals >0 indicate that climate change increases trait values (e.g., climate change factors increase food consumption in most analyses), and negative effect sizes indicate that climate change factors reduce trait values (e.g., temperature reduces developmental time in panels A and D). If the 95% confidence intervals span 0, the meta-analysis did not detect a significant effect of a treatment on a trait. Only one study manipulated springtime freezing conditions (vernal freeze), so this result must be treated with caution. Not all treatment by trait combinations are reflected in each of the subsets of data (panels B-D).

Elevated temperatures significantly accelerated herbivore developmental rates (Fig. 3). In analyses of the direct effects of climatic factors on herbivores, increased temperatures depressed growth rates and did not influence consumption (Fig. 3C). Yet, in analyses of the indirect effects of climate change (when only plants were exposed to climate change factors), elevated temperature augmented consumption and growth rates, and accelerated developmental time (Fig. 3D). While we did not examine effects of temperature on plant chemistry, Zvereva and Kozlov (2006) showed that elevated temperature decreased carbohydrates and phenolics, increased terpenoids, and had little effect on leaf C:N ratio. These changes in plant chemistry could reduce food conversion efficiency, prompting a compensatory feeding behavior in insects. Results of our meta-analysis suggest that consumption rates increase not only because of faster development and growth, but also to compensate for depressed host plant nutritional quality under higher temperature.

Our analyses detected increased consumption rates exclusively in agricultural systems and non-native herbivore or plant food species, but not in native ecosystems (Fig. 4). Yet, in native systems, herbivores also had accelerated developmental times and increased growth rates (Fig. 4A). In diverse natural ecosystems, herbivores can feed selectively across hosts (Bernays *et al*., 1994) and optimize foraging efficiency by regulating pre- and post-ingestive nutrient intake (Behmer, 2009). These mechanisms and the polyphagous nature of many herbivores may allow them to compensate for reductions in nutritional quality in certain host species in natural systems (Behmer, 2009). Additionally, reductions in plant nutritional quality may be more likely to arise in agricultural species, which have generally been subject to strong artificial selection for higher nutritional quality (Newell-McGloughlin, 2008; Whitehead *et al*., 2017). Agricultural systems are typically species-poor, restricting the dietary options of herbivores; therefore, increased consumption rates may emerge more frequently in agricultural than in natural systems under climate change. By stimulating herbivore consumption in agricultural systems and herbivore growth and development in natural systems, warmer climates could increase herbivore pressures on plants, as long as temperatures do not exceed performance thresholds (Angilletta *et al*., 2010). We encourage additional empirical studies on native herbivore-plant systems to verify whether accelerated herbivore development and growth rates also increase consumption rates and plant damage

**Fig. 4:**
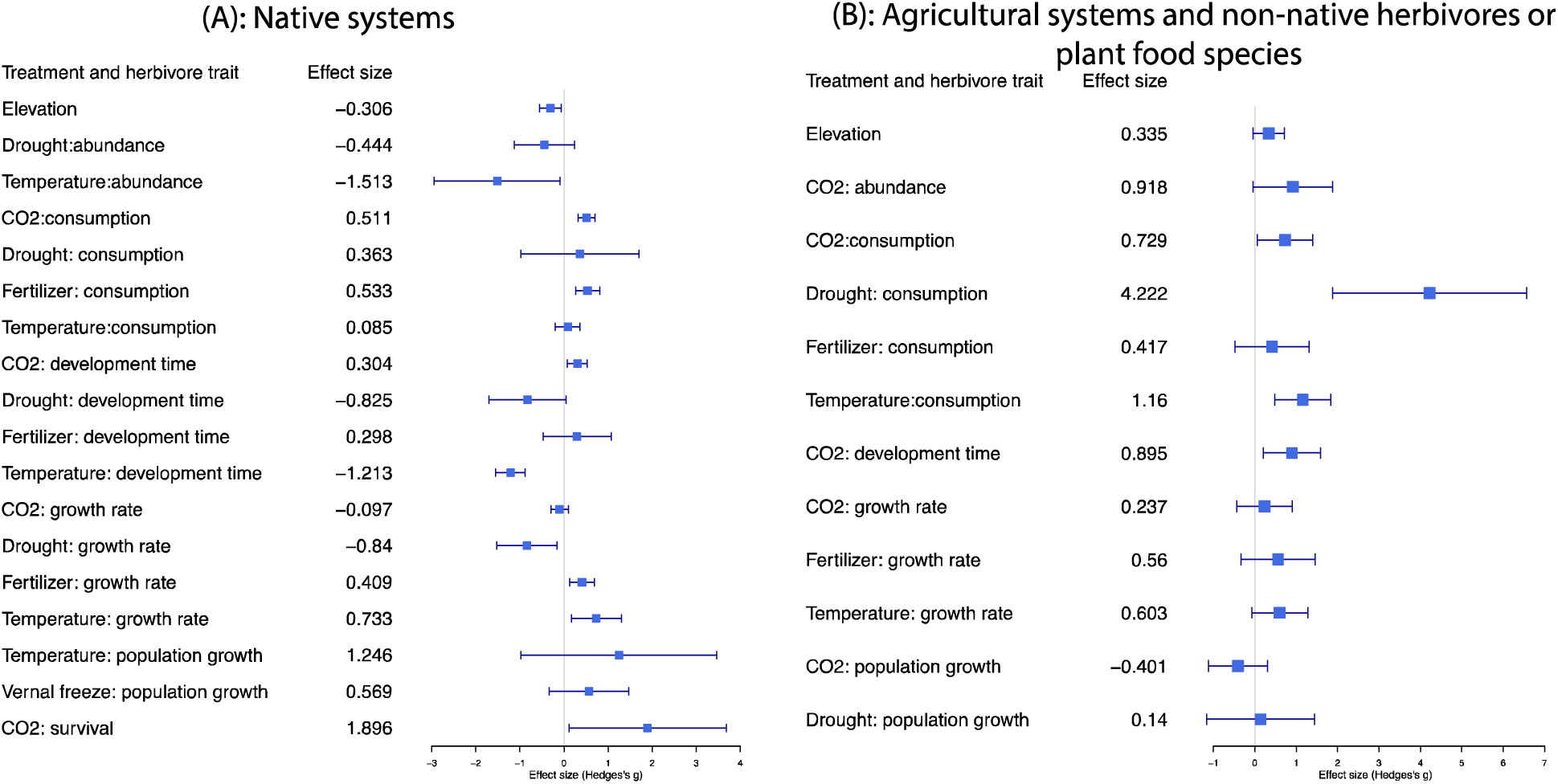
Results of phylogenetically-corrected meta-analysis of herbivore responses at the individual- and population-levels to climate change manipulations in the field and controlled conditions. Plotted are Hedges’s g effect sizes and 95% confidence intervals. We conducted separate meta-analyses for studies conducted in (A) native systems (42 studies), or in (B) agricultural systems and non-native herbivores or plant food species (20 studies).

Elevated CO_2_ increased consumption rates despite delayed insect development and a tendency towards decreased population growth rates (Fig. 3) in both native and agricultural systems (Fig. 4). Since we also identified these patterns in the analysis of plant-mediated indirect effects (Fig. 3D), we conclude that increased consumption rates are likely driven by decreased nutritional value of plant tissues (see also Stiling & Cornelissen, 2007; Robinson *et al*., 2012). Across the six studies of the direct effects of climate change, elevated CO_2_ also increased consumption rates and delayed development, yet accelerated population growth rates (Fig. 3C). However, in half of these studies of direct effects, herbivores were fed artificial diets (Xie *et al*., 2015; Akbar *et al*., 2016; Liu *et al*., 2017), or examined in growth chamber feeding trials (Ipekdal & Caglar, 2012; Lemoine *et al*., 2013; Lemoine *et al*., 2014), and may not be representative of natural field conditions. Effects of elevated CO_2_ and other climatic factors on herbivore performance under field conditions should become a priority of future research. Taken together, our results strengthen the general conclusion that elevated CO_2_ levels affect insect performance through plant-mediated mechanisms (Pincebourde *et al*., 2017). However, herbivore compensatory feeding behavior does not seem sufficient to counteract declines in leaf nutrient quality, and insect performance and population growth rates may suffer under elevated CO_2_.

Drought stress augmented herbivore consumption in agricultural – but not natural – systems (Fig. 3 and 4), but did not affect insect abundance, developmental time, growth rates, or population growth rates in either agricultural or natural settings (Fig. 3). The effects of drought on insects depend on the length and severity of drought episodes, and on the insect feeding guilds (Huberty & Denno, 2004; Gely *et al*., 2019). Our meta-analysis is dominated by Lepidoptera, mainly represented by leaf chewers. For these species, increased consumption rates under drought does not appear to translate into elevated individual or population growth rates. For instance, drought depressed larval growth rates in *Hyalophora cecropia* (Lepidopetra) (Scriber, 1977), and survival and fecundity in *Neodiprion gillettei* (Hymenoptera) (Mcmillin & Wagner, 1995). As for other feeding guilds, severe drought reduces plant turgor, which can restrict the availability of nitrogen-containing compounds, limiting performance of sap-feeders (Gely *et al*., 2019). Additionally, drought can induce greater leaf toughness (Wright & Westoby, 2002), which could reduce palatability for herbivores, and can increase the concentrations of chemical defenses, which could deter herbivores (Gely *et al*., 2019). In our meta-analysis, increased consumption rate under drought was entirely driven by studies that exposed plants but not herbivores to drought (Fig. 3D). Thus, it seems more likely that plants become less resistant to herbivores under drought.

Nutrient fertilization increased consumption rates and tended to enhance herbivore growth rates (Fig. 3) in native but not agricultural systems (Fig. 4). Fertilization is already commonly used in agricultural systems. However, in natural systems, increasing N deposition has been accelerated in many parts of the world, which may change plant-herbivore interactions (Hattenschwiler & Schafellner, 1999). Under intensified N-fertilization, plants have enhanced growth and increased protein concentrations in their tissue, which could increase susceptibility to herbivores (Henn & Schopf, 2001). Congruent with expectations, fertilization enhances insect performance via plant-mediated effects (Fig. 3D), but these indirect effects may be counteracted by elevated CO_2_. For example, the negative CO_2_ effects were greater than the positive N effects on *Lymantria monacha* larval performance (Hattenschwiler & Schafellner, 1999). Our results indicate that increased fertilization may enhance herbivore performance, particularly in natural systems where nutrient limitation may have been important historically.

Our analyses of interactive effects of climate change factors on herbivore performance and foraging behavior were restricted to 16 studies that evaluated interactions between CO_2_ and drought (n= 2 studies), CO_2_ and temperature (n= 8 studies), CO_2_ and fertilizer (n= 5 studies), and temperature and drought stress (n=1 study) (Fig. S5). The overall meta-analysis captures additive effects of these climatic factors from these studies, but does not evaluate whether these factors interact to dampen or exaggerate herbivore responses. We conducted a complementary analysis to evaluate synergistic effects using multifactorial studies only (following Gurevitch *et al*., 2000); however, we failed to detect interactive effects because of the limited numbers of studies and their highly species-specific results. For example, Zhang *et al*. (2018) showed that the combination of elevated temperatures and CO_2_ decreased growth rate and food conversion efficiency for *Spodoptera litura*, an agricultural pest particularly destructive for soybean. However, studies conducted with multiple herbivore species, feeding on several host species, often reported contrasting results. Williams *et al*. (2000) detected no interactive effects of CO_2_ and temperatures on gypsy moth (*Lymantria dispar*) performance fed on sugar maple (*Acer saccharum*), but the negative effects of elevated CO_2_ for herbivore performance were slightly dampened under elevated temperatures when gypsy moth feed on red maple (*Acer rubrum*). Similarly, Johns *et al*. (2003) detected increased feeding under elevated CO_2_ and temperatures only in one of two chrysomelid beetles. Finally, other studies failed to detect any interactive effects between climate change factors (Veteli *et al*., 2002; Himanen *et al*., 2009; Niziolek *et al*., 2013; Gherlenda *et al*., 2015). We suggest the differences in patterns observed by these empirical studies stem from varying experimental designs, where herbivores and/or plants were under climate factor manipulations, and differences study systems or settings.

### Plant dataset

Our meta-analysis revealed that some plants may suffer more damage from herbivory under climate change (Table S8-S11). For one, floral herbivory increased under elevated temperature and drought, yet this result is based on only two studies conducted on native perennial plants (Fig. 5A, B), and should be interpreted cautiously. We observed significantly increased leaf damage for annual plants under elevated CO_2_, temperature and drought, which is likely driven by dynamics in natural – not agricultural – ecosystems (Fig. 5C). We observed no overall change in leaf damage for perennial plants under climate change factors. Annual plants reproduce during a single growing season, and the developmental switch from vegetative growth to reproductive growth occurs early in their life-cycle. Therefore, annuals often have a higher investment in reproductive structures compared to vegetative growth than perennials (Bazzaz *et al*., 1987), and may not be able to compensate for herbivore damage as effectively as perennials, which continuously invest in growth, even after reproduction. Additionally, perennials may allocate more resources to physical and chemical herbivore defenses than annuals (Bazzaz *et al*., 1987). Additional studies in both perennial and annual native species are needed to evaluate how climate change factors influence above- and belowground herbivory.

**Fig. 5:**
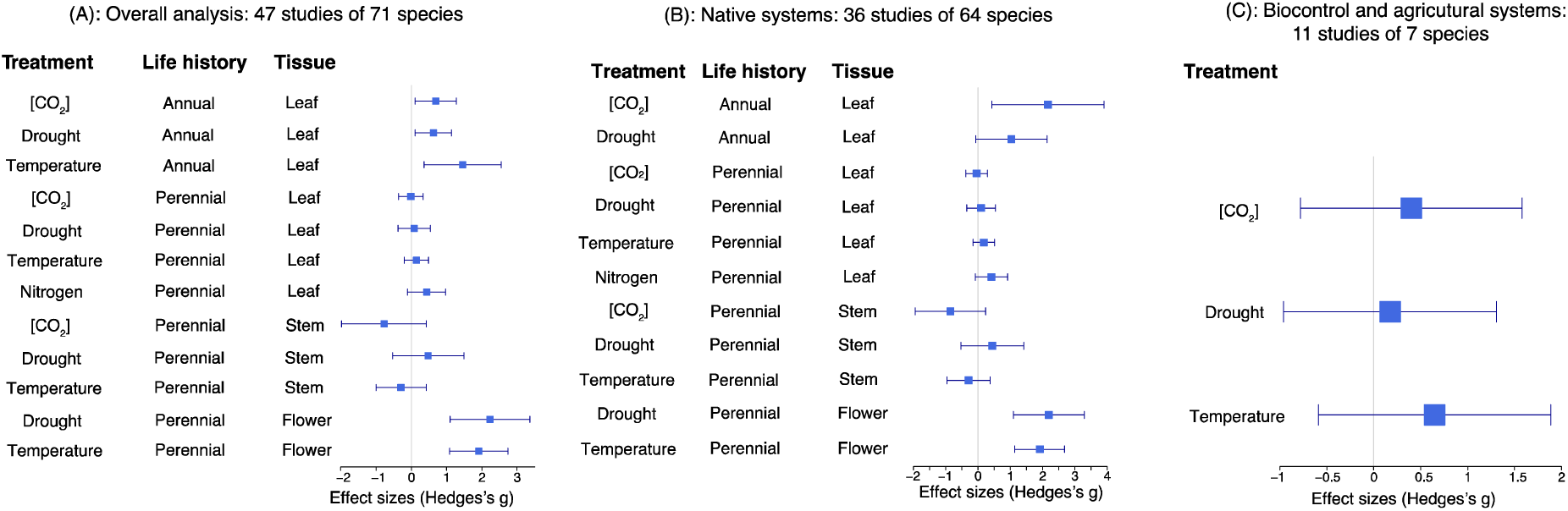
Results of phylogenetically-corrected meta-analysis of herbivore-induced plant damage to climate change manipulations in the field and controlled conditions. Plotted are Hedges’s g effect sizes and 95% confidence intervals. We present the results for separate meta-analyses including (A) the full dataset including all 47 studies, and studies conducted in (B) native systems, or in (C) biocontrol and agricultural systems. As studies differed in design, the treatments were not all applied to each life history or tissue type category. Positive values with 95% confidence intervals >0 indicate that climate change increases trait values, and negative effect sizes indicate that climate change factors reduce trait values. If the 95% confidence intervals span 0, the meta-analysis did not detect a significant effect of a treatment on a trait. Only two studies examined floral herbivory, so these results must be treated with caution.

Our meta-analysis included only 6 studies that examined interactive effects of CO_2_ and temperature (n = 2 studies), temperature and drought (n = 2 studies), temperature and nitrogen fertilization (n = 1 study), and CO_2_ and drought (n = 1 study) (Fig. S8). Therefore, we were unable to analyze interactive effects of climate change factors on plant damage that extended beyond additive effects of each climate change factor. Both studies that simultaneously examined CO_2_ and temperature found no interactive effect on feeding damage (Johns *et al*., 2003; Himanen *et al*., 2009). Yet, increased herbivore damage was found under interactive effects of temperature and drought on St. John’s wort flowers (Fox *et al*., 1999) and red oak leaves (Rodgers *et al*., 2018). In addition, Lu *et al*. (2015) found greater rates of root galling under elevated temperature and nitrogen.

### Taxonomic and geographic breadth of plant and herbivore datasets

Our plant dataset included 71 angiosperm species, spanning 52 genera and 27 plant families, most represented by Fabaceae and Fagaceae (Fig. S7a). In our meta-analysis, 90% of the studies examined perennials, 54% examined herbaceous plants, and 90% of studies quantified foliar herbivory (Fig. 7b, c, d). Woody plants represent ∼45-48% of species globally (FitzJohn *et al*., 2014), and perennials represent about 60% of all seed plants and 40% of domesticated species (Miller & Gross, 2011, Miller pers. comm.), illustrating that herbaceous and perennial species were likely overrepresented in our datasets. More studies should focus on plant-herbivore interactions in annual plants, alternative life-forms, especially shrubs, and other tissues, such as roots (Johnson *et al*., 2016), and reproductive structures.

The taxonomic breadth of our herbivore dataset included 57 species, spanning 34 families and 8 insect orders, most represented by Lepidopterans (Fig. S4a). This bias towards Lepidopterans and the leaf-chewing feeding guilds was also documented in earlier reviews (Bezemer & Jones, 1998; Stiling & Cornelissen, 2007). Are plant-herbivore studies taxonomically biased or is the dominance of Lepidopterans representative of insect herbivore diversity? Lepidopterans are among the three most diverse insect orders (Goldstein, 2017). Although Coleoptera and Hymenoptera are more diverse insect orders, 100% of Lepidopterans are herbivorous species, while only 26% and 7% of Coleoptera and Hymenoptera are herbivores, respectively (Wiens *et al*., 2015). Thus, empirical studies likely are not unduly biased toward Lepidopterans.

We found another striking bias in both herbivore and plant datasets: the vast majority of studies were conducted in the Global North. Only 5 studies examined plant damage and herbivore performance in southern latitudes, illustrating a clear underrepresentation of plant-insect interactions in the Global South. We call for funding to support future studies on the consequences of climate change for plant-insect interactions in the Global South, especially in tropical biodiversity hotspots.

### Future directions

Our herbivore meta-analysis revealed that studies are not always performed with herbivores and plants both experiencing climate change manipulations. While having only plants or herbivores under manipulated climate change factors can help disentangle direct and plant-mediated effects of climatic factors, we suggest that the most realistic results would come from exposing both plants and herbivores to changing environmental conditions.

More studies using multifactorial designs are urgently needed to achieve a realistic understanding of how current climate change influence plant-herbivore interactions. The joint effects of climate change factors likely depend on the magnitude of changes, on herbivore feeding guilds, and on species-specific interactions, and climate change factors may interact in complex ways (Jamieson *et al*., 2017). For example, forest tent caterpillar (*Malacosoma disstria*) consumption increased on aspen under drought stress, regardless of CO_2_ levels, but declined more strongly on drought stressed maple leaves under elevated CO_2_ (Roth *et al*., 1997). Similarly, population growth rates of two-spotted spider mite (*Tetranychus urticae*) increased only when elevated CO_2_ was combined with moderate drought stress (Sinaie *et al*., 2019). Additionally, temperature affected the magnitude and the direction of plant and herbivore responses to elevated CO_2_ in three chrysomelid beetle species and two plant species (Veteli *et al*., 2002; Johns *et al*., 2003). Furthermore, increasing temperature and CO_2_ may have opposing effects on herbivore performance. We strongly encourage future multifactorial studies to evaluate plant and herbivore responses to the complex suite of climatic conditions that are changing simultaneously, ideally under field conditions that capture natural variation in numerous biotic and abiotic conditions simultaneously (Körner, 2003; Moles *et al*., 2011; Rasmann *et al*., 2014b). While we are certainly not the first to call for multifactorial studies (Bale *et al*., 2002; Massad & Dyer, 2010; Giron *et al*., 2018; Hartley & Beale, 2019), our literature search revealed a dearth of multifactorial studies. Climate change factors could operate independently, synergistically or antagonistically, and multifactorial experiments are needed to generate robust predictions about plant-herbivore interactions under simultaneous changes in CO_2_, temperature, precipitation and other variables.

## III. Evolutionary consequences of climate change for plant-herbivore interactions

One fundamental question is whether species will be able to adapt fast enough to track rapid environmental change (Visser, 2008). Plants have evolved a variety of chemical and morphological traits that allow them to resist or tolerate herbivores, and insects have evolved traits that allow them to overcome many plant defenses (Ratzka *et al*., 2002; Glauser *et al*., 2011; Jander, 2014; War *et al*., 2018). The evolution of plant and insect traits related to herbivory and defense can happen rapidly enough to influence ecological dynamics. For example, *Oenothera biennis* plants protected from herbivory evolved reduced resistance and increased competitive ability over the course of a 4-yr experiment (Agrawal *et al*., 2012). Additionally *Brassica rapa* plants evolved rapidly in response to pollinators and foliar herbivores (Ramos & Schiestl, 2019). Furthermore, heterogeneity in herbivore abundance across the landscape can influence genetic variation in plant defenses. For example, geographic variation in the defense locus *GS-ELONG* in the model plant *Arabidopsis thaliana* is associated with aphid abundance (Züst *et al*., 2012). Insect species often have fast developmental rates, short generation times, and high reproductive rates, which can lead to rapid evolution to novel conditions, like pesticides (Hawkins *et al*., 2019) and introduced plant hosts (Carroll *et al*., 2005). Evolutionary changes in herbivorous insects influence host preferences (Singer & Thomas, 1996; Thomas *et al*., 2001) and herbivore distributional shifts (Haag *et al*., 2005).

Adaptive responses to climate change depend on the magnitude of novel selection and the degree of genetic variation in functional traits. These quantitative genetic parameters are hard enough to quantify in response to changing abiotic conditions (Etterson & Shaw, 2001; Bemmels & Anderson, 2019; Torres-MartÍnez *et al*., 2019). The situation becomes even more challenging when interacting species also impose strong selection on each other, and when one partner (often the insect herbivore) has a much faster generation time than the other (often the plant host). Is existing genetic variation within plant and herbivore populations sufficient to adapt to ongoing climate change in the long-term? Will rapid contemporary evolution of arthropod herbivores further depress the fitness of native plant species? Could gene flow across plant populations accelerate adaptation if gene flow occurs primarily from populations that evolved with a diverse and abundant herbivore community into populations that historically experienced a more depauperate community? Could adaptive responses to climate change in plants or herbivores be constrained by increased herbivory or defenses, respectively? Our meta-analysis suggests that climate change likely exerts strong selection on plant and herbivore traits, but we know very little about the longer-term evolutionary consequences of ongoing environmental change for plant-herbivore interactions.

Anthropogenic environmental changes such as climatic change, habitat fragmentation, pollution and urbanization likely interact to influence the co-evolutionary dynamics of plants and herbivores (Leimu *et al*., 2012; Miles *et al*., 2019). Changes in temperature can influence plant chemistry and phenology, as well as insect growth and feeding rates (Bale *et al*., 2002; Huberty & Denno, 2004; Zvereva & Kozlov, 2006). Because of the complex interactions involved, the long-term consequences of these environmental changes on community dynamics are likely to be difficult to predict. Our review has focused primarily on arthropod herbivores because fewer studies have evaluated changing plant-herbivore interactions under climate change for mammalian herbivores or other taxonomic groups (but see Brodie *et al*., 2012; Choi *et al*., 2019). Nevertheless, mammalian populations are declining globally (Collen *et al*., 2008; Harris *et al*., 2009; Ripple *et al*., 2015), which could decrease plant damage from larger herbivores. Future studies using a variety of complementary approaches will be needed to predict how environmental changes will influence plant-herbivore eco-evolutionary dynamics.

Traditional approaches to investigating plant-herbivore coevolution have used techniques such as phylogenetic analysis (Ehrlich & Raven, 1964). However, plant-herbivore eco-evolutionary dynamics can occur rapidly and can also be studied via lab and field experiments (Agrawal *et al*., 2012; Züst *et al*., 2012; Ramos & Schiestl, 2019). Another promising technique is the resurrection approach of comparing ancestors and descendants under common conditions to directly examine evolutionary change (Franks *et al*., 2018b). One study used ancestral and descendant seeds of rapid-cycling *Brassica rapa* plants and showed that the evolutionary changes that occurred through artificial selection for rapid cycling resulted in changes to herbivore preference and performance (Franks *et al*., 2018a). Resurrection studies may be more challenging to use for insects than plants, as seeds can often be stored long-term. However, for certain herbivore species, dormant eggs may be retrievable from the soil bank, or frozen and revived for comparison with contemporary generations (Kerfoot & Weider, 2004; Franks *et al*., 2018b). Alternatively, lab colonies may be maintained over generations and serve as a link to the past (Cooper *et al*., 2003). Similarly, in experimental evolution studies, colonies could be reared on artificial diets to eliminate selection by plant traits.

Resurrection studies and experimental or artificial selection experiments can be combined with genomics to study adaptation from standing genetic variation (Schlötterer *et al*., 2015). Genomic studies can examine how spatially variable selection, mediated by plant-herbivore interactions, can maintain genetic variation within natural populations of host plants and herbivores (Gloss *et al*., 2013). For example, geographic variation in herbivory can drive adaptive evolution and maintenance of polymorphism in plant defense genes (Prasad *et al*., 2012), while spatial mosaics of host plants may also maintain phenotypic variation in herbivores (Kant *et al*., 2008). Experimental evolution studies (Ramos & Schiestl, 2019), studies using genome-wide sequencing (Gloss *et al*., 2016), and studies that involve experimental manipulations of factors that are being altered with anthropogenic environmental change (Leimu *et al*., 2012) will all be useful for helping to understand and predict changes in dynamic interactions between plants and herbivores.

### Conclusions

Our meta-analysis revealed that climate change factors can increase herbivore consumption rates, likely leading to greater foliar damage to annual plants and floral damage to perennial plants. Furthermore, we found that increased CO_2_ concentrations delayed the development of arthropod herbivores, whereas increased temperatures accelerated development. We hypothesize that some insect herbivores may shift from one to multiple generations per year under climate change. We caution that these results focus almost entirely on interactions between arthropod herbivores and foliar tissues in temperate systems, and that most studies manipulated a single climatic condition at a time. We encourage future studies that evaluate additional plant-herbivore dynamics under climate change, including florivory and seed predation, both of which are tightly associated with plant fecundity (Parachnowitsch & Caruso, 2008). Future studies will reveal how climate change will influence belowground interactions and plant-soil feedback responses (Bezemer *et al*., 2013; Lu *et al*., 2015; Ourry *et al*., 2018). Finally, several studies have evaluated mammalian herbivory under climate change (Brodie *et al*., 2012), and future experiments that disentangle the contributions of multiple taxonomic groups to increased herbivory under climate change will increase our capacity to predict plant and herbivore population persistence. Given the strong immediate effects of climate change on plant and herbivore functional traits, novel conditions could impose strong selection and alter long-term evolutionary dynamics.

## ACKNOWLEDGEMENTS

We thank Tom Pendergast for discussions about plant-herbivore interactions and Rachel MacTavish, Derek Denney, Mia Rochford, Kelly McCrum, and Chazz Jordan for comments on a previous draft. The Anderson and Franks labs are supported by the U.S. National Science Foundation (Anderson: DEB-1553408 and DEB-1655732; Franks: IOS-1546218). We thank L. Delph and two anonymous reviewers for providing constructive criticism.

## AUTHOR CONTRIBUTIONS

E.H. conducted the literature search, extracted data, and wrote the manuscript. C.B. contributed to data extractions and commented on the manuscript. S.F. contributed to manuscript writing and editing. I.J. contributed to the introduction and commented on the manuscript. J.T.A. developed the research framework, extracted data, conducted the meta-analyses, and wrote the manuscript.

